# Classifying smartphone-based accelerometer data to obtain validated measures of subject activity status, step count, and gait speed

**DOI:** 10.1101/160317

**Authors:** Yong Jun Kwon, Thawda Aung, Sarah M Synovec, Anthony D Oberle, Cassia Rye Hanton, Jackie Whittington, Evan H Goulding, Bradley C Witbrodt, Stephen J Bonasera, A Katrin Schenk

## Abstract

**Background:** The ubiquitous spread of smartphone technology throughout global societies offers an unprecedented opportunity to ethically obtain long-term, highly accurate measurements of individual physical activity. For example, the smartphone intrinsic 3-D accelerometer can be queried during normal phone operation to save time series of acceleration magnitudes (in each of the component directions) for near-real time or post processing.

**Objective:** We describe simple, straightforward algorithms (based on windowed Fourier analysis) for accelerometer data quality control and behavioral classification.

**Methods:** To maximize the clinical utility of our classifications, we focused on differentiating the following conditions: forgotten phone, subject resting, low physical activity, high physical activity. We further differentiated high physical activity into epochs of walking and climbing stairs, and further quantified walking to infer step count and gait speed.

**Results:** We validated these algorithms in 75 individuals, in both laboratory (treadmill) and naturalistic settings. Our algorithm performance was quite satisfactory, with accuracies of 92-99% for all behavioral categories, and 87-90% for gait metrics in naturalistic settings.

**Conclusions:** We conclude that smartphones are valid and accurate platforms for measuring day-to-day physical activity in ambulatory, community dwelling individuals.

## Introduction

Decades of advances in the design and manufacture of integrated circuits, combined with decreases in the cost of these components, are fueling an ongoing revolution in the capacity to make continuous real or near-real time measurements and analyses of dynamic processes. These kinds of approaches are particularly well-suited for studying the complex mixture of periodic, nonperiodic, and stochastic events that comprise animal or human behavior. Better understanding of behavioral structure is highly relevant in many diverse fields such as migratory ecology [1], law enforcement and surveillance [2], robotics [3], economics [4], and behavioral neuroscience [5].

Human physical activity is a clinically and economically important behavior that is particularly well suited for continuous measurement. Consequences of physical inactivity can include significant and costly chronic health problems such as obesity, type II diabetes, vascular disease, hypertension, colon cancer, and depression [6]. Increased physical activity can help prevent these illnesses, improve bone, joint, and muscle function, and even decrease the risk of premature death. However, most tools validated to assess individual physical activity provide a partial portrayal of overall status. For example, validated surveys (*e.g.* PROMIS, BRFSS, IPAQ; [7–9]), while highly useful as clinical tools to stratify physical activity in community-dwelling populations, are not useful for providing day-to-day guidance on overall activity. Similarly, clinical physical performance batteries [*e.g.* 10] provide a useful, quick clinical screen for persons at risk for functional loss, but are relatively insensitive to smaller improvements or losses of physical function. New measures of physical function that reflect daily performance must ultimately be developed in order to meet the needs of community-dwelling individuals seeking to increase and maintain physical activity for wellness or rehabilitation.

Inferring physical activity from continuous measurements (such as those from an accelerometer) is not a trivial task, and an extensive literature has risen to describe this problem [for review, 11]. Many of these machine-learning based classification algorithms have been proven to demonstrate significant accuracy in discriminating between different modes of physical activity [12]. However, many of these approaches require significant computational efforts that are not feasible for near-real time interventions (particularly in rural or underdeveloped areas with unreliable access to telecommunication infrastructure). Furthermore, clinically-relevant metrics – particularly with regard to an aging population – such as gait speed, number of footsteps, and total activity duration, could be immediately accessible on a local device such as a cell phone. Far fewer algorithms have been developed for this situation [13]. Finally, selection of the appropriate sensor(s) for measuring physical activity in community-dwelling populations is also not an insignificant consideration. Most measurements of physical activity ultimately rely on the subject wearing some kind of accelerometer. There are many commercial providers of accelerometer technology with costs ranging from tens to thousands of dollars (US) per subject. However, with the exception of devices worn as watches or integrated into clothing, many of these devices require active effort on the part of the subject’s day-to-day life in order to facilitate data collection. Of note, advances in microelectromechanical systems have markedly shrunk the cost and size of accelerometer technology. Inexpensive indwelling accelerometers are now components of many commonly used devices, including automobiles (braking and airbag control), games (Wii remote), sports training (footpods), and personal electronics (computers, cell phones).

Here, we describe our first efforts to validate algorithms that extract clinically-relevant features of activity (gait speed, number of footfalls, and activity duration) in a near-real time, continuous, noninvasive manner in community-dwelling, ambulatory, older adults. To maximize subject ease-of-use, we measured physical activity using the sensor capabilities of inexpensive “smartphones.” We describe our algorithms for data conditioning, as well as determination of (1) whether the phone was on the subject’s person, (2) subject overall activity/inactivity, (3) gait speed, and (4) step count. We show that these algorithms closely match gold standard measures of these outcomes. We conclude that our previously-described approach for measuring functional status in community-dwelling adults [14] provides valid outcome measures of activity and gait.

## Methods

*Validation samples.* Subject datasets to develop and validate the below-described algorithms were obtained from three separate sources. For step count and gait speed determination algorithms, we used data from a prior clinical study examining gait under the highly controlled conditions of treadmill walking [15]. This study provided acceleration/step count/gait speed characteristics from 17 young (19-35 y/o), 19 middle-aged (36-65 y/o), and 19 older (>65 y/o) individuals. Subjects alternated 5 minute periods of gait at defined treadmill speeds (between 0.4 and 7 mi/hr, depending upon subject tolerance and fitness) with 1 minute rest periods.

Data for development and validation of algorithms examining activity, step count, and gait speed in naturalistic environments was collected by two separate young subjects (ADO, SMS). To validate gait speed in a naturalistic environment, they walked on an outdoor track of known length (0.25 mi) while using a metronome to keep constant pace. Data collected between gait speeds of 2-5 mph were used for further analysis to mimic adult regular walking speeds. Additionally, these subjects performed numerous activities while wearing the cell phone, including sightseeing, shopping, going to the movies, napping, sleeping, and taking auto trips. The subjects kept extensive, one-minute resolution behavioral diaries documenting their activities during this time. This data was used to develop and validate our behavioral classification algorithms. All in all, these subjects collected more than 3485 minutes of behavioral data, including 466 minutes of “phone forgotten,” 1150 min of low physical activity (including 684 min of resting or sleeping), 1564 min of driving, and 771 min of high physical activity (including 619 minutes walking and 152 min climbing stairs). Half of this data was used for model development, and the other half used for model validation.

Finally, we applied the above-validated algorithms to cell-phone measured activity patterns obtained over a full day from a group of 18 normal adults. Inclusion characteristics included age > 55, independent living in the community, and the absence of significant or uncontrolled medical or psychiatric problems. All participants had no evidence of cognitive impairment (MOCA>24, Montreal Cognitive Assessment Test, [16]). Many subjects were enrollees in UNMC Engage, an exercise- and community activity program offered by the UNMC Home Instead Center for Successful Aging. We obtained 354 hours of activity data from this group, with an average contribution of 19.18 hours, a maximum contribution of 20.74 hours, and a minimum contribution of 15.27 hours.

All studies were performed in complete accordance and approval of the UNMC Institutional Review Board (IRB).

*Protocol.* All activity data was collected using a Nokia N79 cellular phone (White Plains, NY). The N79 has a built-in accelerometer with a dynamic range of ±2 g in the three axes (*e.g.*, x, y, z). The accelerometer was sampled at approximately 7-8 Hz using the PyS60 *sensor* framework running on the Symbian S60 FP2Ver3 operating system. Raw acceleration data was written to the phone flash drive. Subjects kept the phone in either hip pocket and simultaneously wore an electronic pedometer (New-Lifestyles NL-2000; Lees Summit MO) on their belt to estimate total footsteps.

*Approach to activity data classification.* We focused on activity metrics with clinical relevance, high face validity, and proven linkage to important patient outcomes that could easily be implemented for translational research. Figure 1 depicts our overall strategy. First, we perform quality control (QC) to condition accelerometer data for classification. Following data QC, we classify accelerometer data into epochs of “forgotten” versus “carried” phone. We classify epochs of “carried” phone into periods of low physical activity (“IPA”) and high physical activity (“hPA”). Epochs of lPA are further classified into periods of driving (“driving”) and periods of other low activity behaviors, such as resting, watching entertainment, *etc*. (“other”). Epochs of hPA are further classified into periods of walking (“walking”) and climbing stairs (“stairs”). Finally, we subclassify walking to determine step count and gait speed, two metrics strongly associated with many important health care outcomes [17–19].

**Figure 1.**
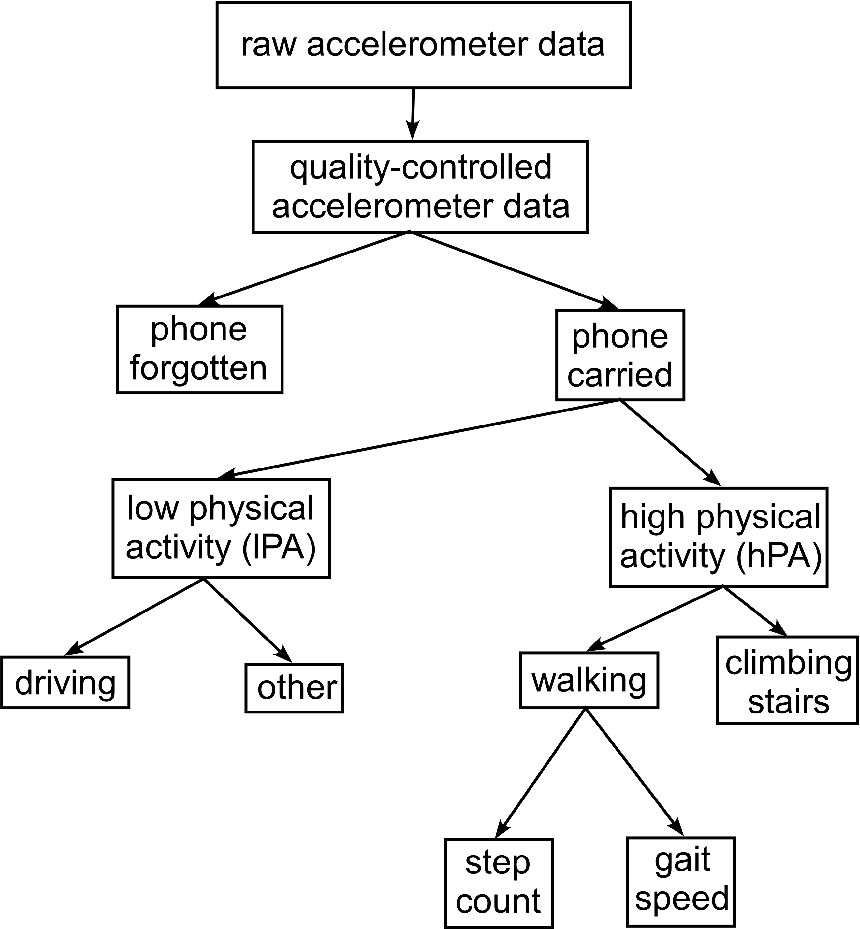
Block diagram depicting overall data quality control/classification approach.

*Accelerometer data quality control.* The Nokia N79 intrinsic accelerometer is not highly precise. For example, when two phones lie motionless on a table, they yield slightly differing acceleration values despite being under the same force (gravity). Moreover, the cell phones output slightly different gravitational acceleration values depending on device relative orientation. Finally, stationary cell phones randomly demonstrate occasional small increases or decreases in baseline acceleration magnitude. These device characteristics affect threshold-based identification techniques used in the physical activity, step count, and gait speed algorithms.

To condition the accelerometer data, we use bandpass filter, window division and acceleration magnitude resetting. First, the acceleration magnitude goes through a butterworth bandpass filter with frequency band of 0.2-0.7 Hz. Acceleration magnitude is then divided into 68 s windows, such that each window contains 512 acceleration points for Fast Fourier Transform computation. We calculate average acceleration magnitude for each window. The window size is wide enough (in the time dimension) so that mean acceleration should approximate gravitational acceleration regardless of subject activity. Then, for each acceleration magnitude point contained in the window, a uniform value is added/subtracted to adjust the window mean acceleration magnitude to 1g. After this procedure, all windows have a mean acceleration magnitude equal to gravitational acceleration without any visible discontinuities (Figure 2).

**Figure 2.**
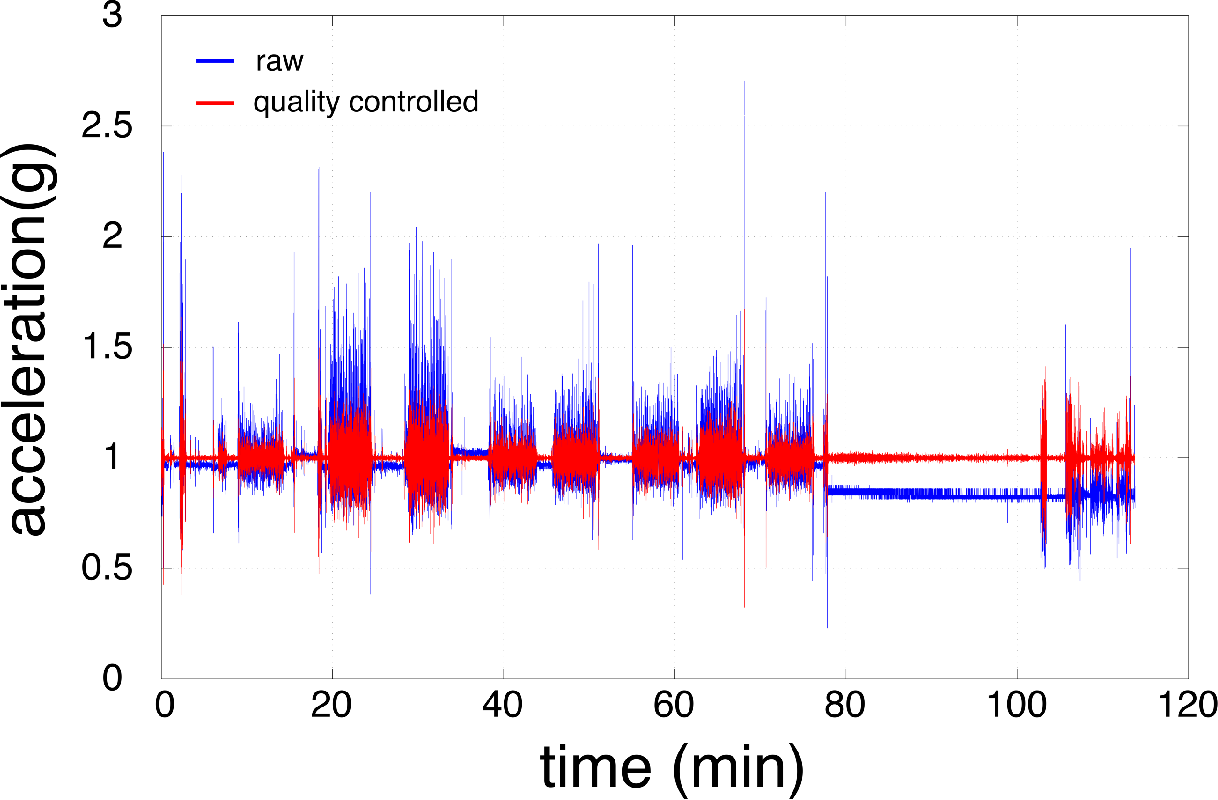
Accelerometer magnitude data quality control. Nearly two hours of representative raw accelerometer magnitude data (blue trace) depicting sudden shifts in baseline accelerometer output, accompanied by corresponding quality-controlled accelerometer magnitude data (red trace). Note that the process of bandpass filtering, window division and acceleration magnitude resetting lead to a continuous accelerometer signal whose magnitude over any epoch averages to 1g.

*Classification of “forgotten” versus “carried” phone.* We calculate the acceleration magnitude root mean square (RMS) over 68 s continuous, nonoverlapping windows. If the window RMS value exceeds an empirically derived threshold (0.0094 g), then the entire window is classified as “carried.” For windows classified in this manner as “forgotten”, we perform a second “pass” analysis and test each acceleration magnitude within the window to determine if it exceeds a second, empirically-determined threshold (1.015 g). Windows initially classified as “forgotten” that have one or more acceleration magnitudes exceeding 1.015 g are reclassified as “carried.”

*Physical activity classification.* Data from “carried” phones having acceleration magnitude RMS ≤0.09375 g (empirically determined) are classified as having low physical activity. Conversely, data from “carried” phone having acceleration magnitude RMS > 0.09375 g are classified as having high physical activity. Figure 3a depicts a representative example of this process. For behavioral windows determined to have low physical activity, we further identify periods where the individual is at rest, sitting quietly (“other”) as having 0.0094 g ≤ RMS ≤0.01 g. Similarly, we use threshold of 0.01 g ≤ RMS ≤ 0.09375 g to identify periods of driving.

**Figure 3.**
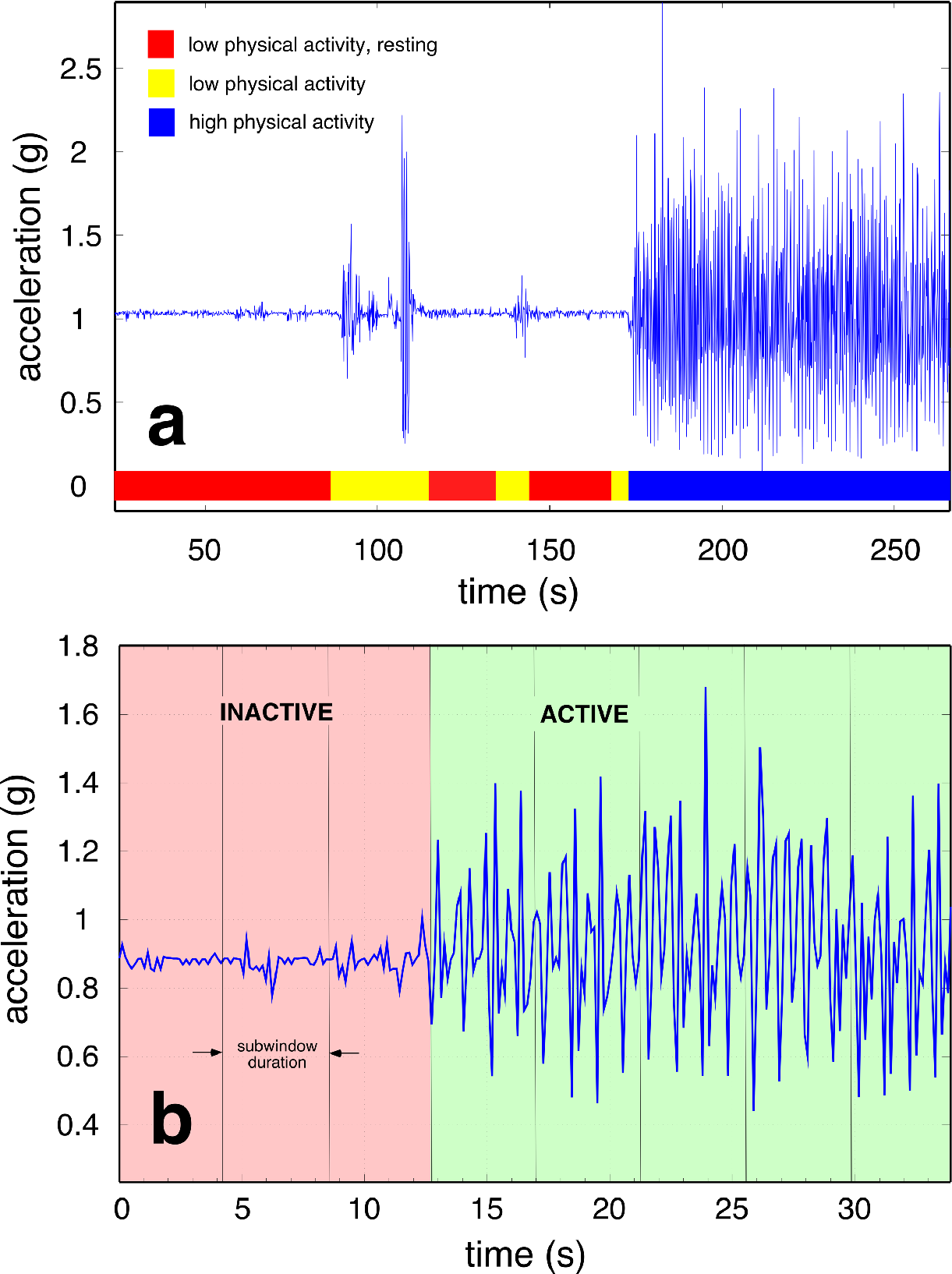
Fourier-based windowing technique for behavioral classification. **a.** Representative quality-controlled acceleration magnitude showing transitions from low physical activity, resting state (red) to low-physical activity, nonresting state (yellow), to high physical activity (blue), **b.** Subwindowing to increase accuracy of inactive/active classification to 4.3 s intervals.

For behavioral windows determined to have high physical activity, we further identify periods where the individual is walking as having thresholds of 0.09375 g ≤ RMS ≤0.15625 g. Similarly, we use threshold RMS values ≥ 0.15625 g to identify periods of climbing stairs. To increase the algorithm precision, hPA epochs were further divided into “sub epochs.” This process allowed us to better determine if specific periods within the initial 68 s window demonstrated low versus high physical activity. Each sub epochs is 4.3 s long. Sub epochs undergo analogous threshold testing to differentiate periods of low and high physical activity. Each 4.3 s window includes 32 data points.

Thresholds were empirically derived from treadmill walking trials, where step counts could be validated against gold standard video for each subject. We chose threshold values that maximized our mean accuracy across all treadmill gait trials.

*Step count determination.* We adapted fast Fourier transforms (FFTs), threshold detection, and sub window techniques to accurately infer subject step count per [20,21]. As above, acceleration magnitude is calculated over 68 second continuous, non-overlapping windows. This window length further assures that periodic signals from walking will be significantly larger than those arising from noise. Subject step frequency was then determined by calculating the FFT across all windows classified as “walking.” Step frequencies greater than expected for walking were reclassified as “other”. After step frequency was determined, the 68 second window is divided into eight subwindows, and each subwindow is thresholded to determine if it includes hPA. From the subwindow that includes hPA, we obtain the proportion of hPA in the original 68 second window. The proportion of walking subwindows was used to determine final window step count, further increasing step count accuracy. The subwindow method is depicted in Figure 3b.

*Gait speed determination.* We estimated individual gait speed for each subject by multiplying gait frequency (derived from the above step count analysis) by the average stride length provided by age- and gender- specific nomograms. For older adult cohort, average stride length was 1.796 feet for women and 2.075 feet for men; for the young and middle-aged cohorts, average stride length was 2.133 feet for men and women [22,23].

*Gold standards for validations.* For data arising from treadmill gait studies, our gold standard for step count was obtained by manual counting of footfalls from video recordings. Gold standard for gait speed was the speed entered on the treadmill. For data lacking video collaboration, we found that pedometer step counts were often unsatisfactory as a gold standard. Pedometer step counts accurately reflected real step counts for our naturalistic activity study involving the two young subjects. However, in the older community living group pedometer step counts and the subject activity log did not agree. We attribute this discrepancy to subjects wearing the pedometer incorrectly. For these situations, we manually reviewed the raw accelerometer data, and were able to identify signal morphologies that identified individual steps within this raw data, and thus obtain an estimated step count. This visually-based method was validated against treadmill data (with gold standard video-based step counts) and found to have 98% accuracy.

## Results

*Creation of automated, reliable behavioral diaries.* We first applied our behavioral classifications to subject one-minute behavioral diaries collected during day-to-day tasks. Comparison of subject journaling versus automated behavioral classification is shown in Figure 4a. Generally, the algorithm detected changes in subject behavior coincidentally with the subject journal. All discrepancies were within one minute onset, and could arise from small time differences between subject time and smartphone time.

**Figure 4.**
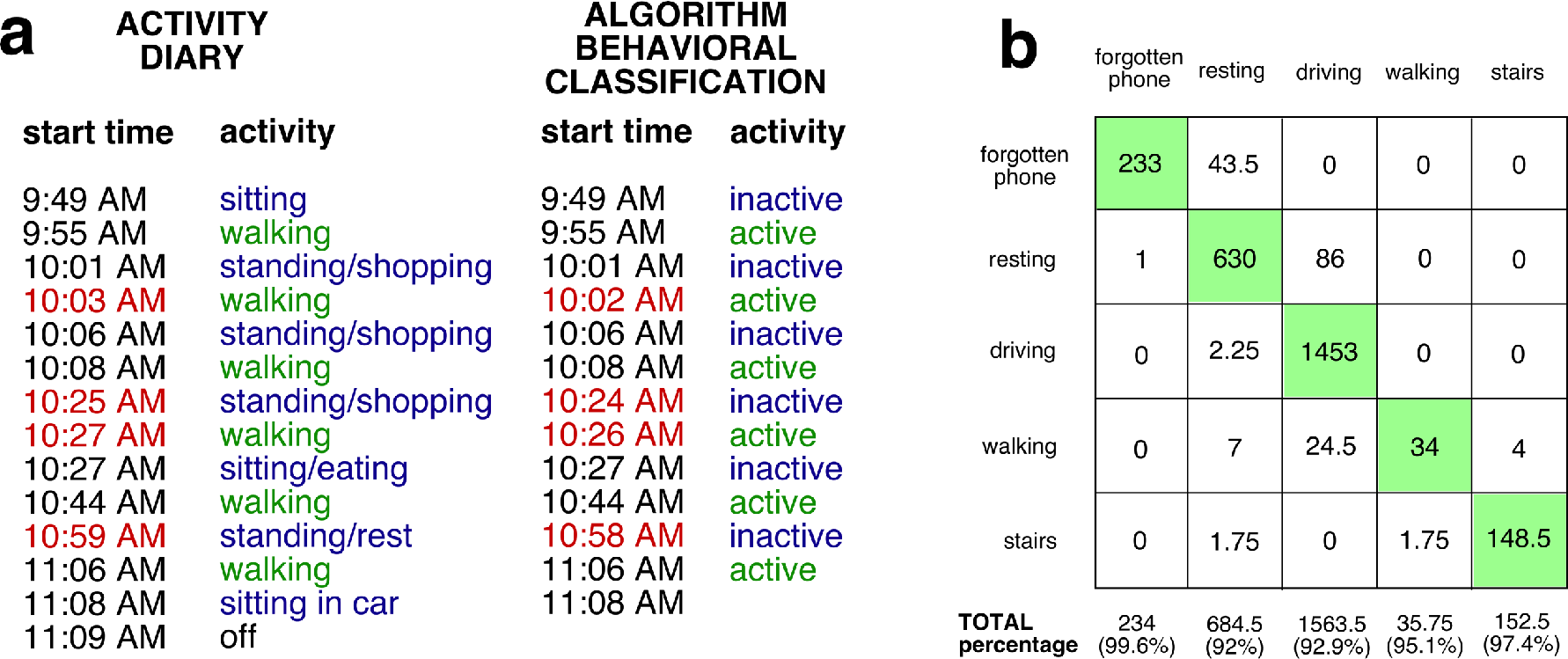
Classification of smartphone acceleration data into clinically-useful activity metrics. **a.** Comparison of subject- vs. smartphone-created behavioral diary. Subject entries (time stamp and behavior) are in the left columns; smartphone classifications (time stamp and behavior) are in the right columns. Note excellent agreement (shown in green) between subject and smartphone with changes in behavioral status (from inactive to active or vice-versa). All discrepancies between subject and smartphone (red) are no longer than one minute in onset. Given the profound difficulty that subjects have performing high-resolution behavioral diaries of their day-to-day lives, this figure suggests that much of this task can be performed more accurately using smartphone data acquisition and behavioral classification, **b.** Confusion matrix depicting the frequency of true positive (along diagonal) and false positive (off diagonal) behavioral classification for forgotten phone, resting, driving, walking, and stairs.

*“Forgotten” and “carried” phone are differentiated with high accuracy.* Our algorithm classified epochs of “forgotten” versus “carried” phone with very high accuracy (99.6%, 234 min classified). The algorithm misclassified one minute of “forgotten” data as occurring in the low physical activity state (confusion matrix provided in Figure 4b). This high specificity suggests that subjects forgetting to carry the phone will not significantly influence our ability to differentiate between low and high physical activity states.

*Low and high physical activity states are differentiated with high accuracy.* In a total of 685 minutes of low physical activity data, 630 minutes (92%) was classified correctly. 43.5 minutes (6.4%) were misclassified as “forgotten” phone. Inevitably, if a subject were completely still, it would be very challenging to differentiate “forgotten” phone from low physical activity. However, low activity states are usually not characterized by a complete absence of movement. We also had a low false positive rate of calling lPA activities as hPA. Specifically, we misclassified 7 min (1%) of IPA activity as walking, and 1.75 min (0.3%) of IPA activity as climbing stairs.

Our thresholding algorithm performed well at differentiating low physical activity states into driving and other. We properly classified 1453 of 1563 min (92.9%) of driving, with 86 min (5.5%) misclassified as other and 24.5 min (1.6%) misclassified as hPA.

Our thresholding algorithm also performed well at differentiating high physical activity states into walking and climbing stairs. We properly classified 34 of 36 min (94.4%) of walking, with 2 min (5.6%) misclassified as climbing stairs. Similarly, we properly classified 148.5 of 153 min (97.4%) of climbing stairs, with 4 min (2.6%) misclassified as walking.

*Accurate quantification of step counts in both laboratory and naturalistic settings.* To first test the validity of our model for step counts, we examined its performance under the highly controlled situation of treadmill locomotion. In all of our figures comparing measured step counts versus predicted step counts, the diagonal line depicts the performance of a perfect model. Points falling above the line suggest that the model finds more steps than were observed; points falling below the line suggest that the model finds fewer steps than were observed. As demonstrated in Figure 5a, we had an overall accuracy of 93% for all age groups (94.5% for young, 93.6% for middle-aged, 94.7% for aged). The model did not consistently over- or under- estimate steps counts.

**Figure 5.**
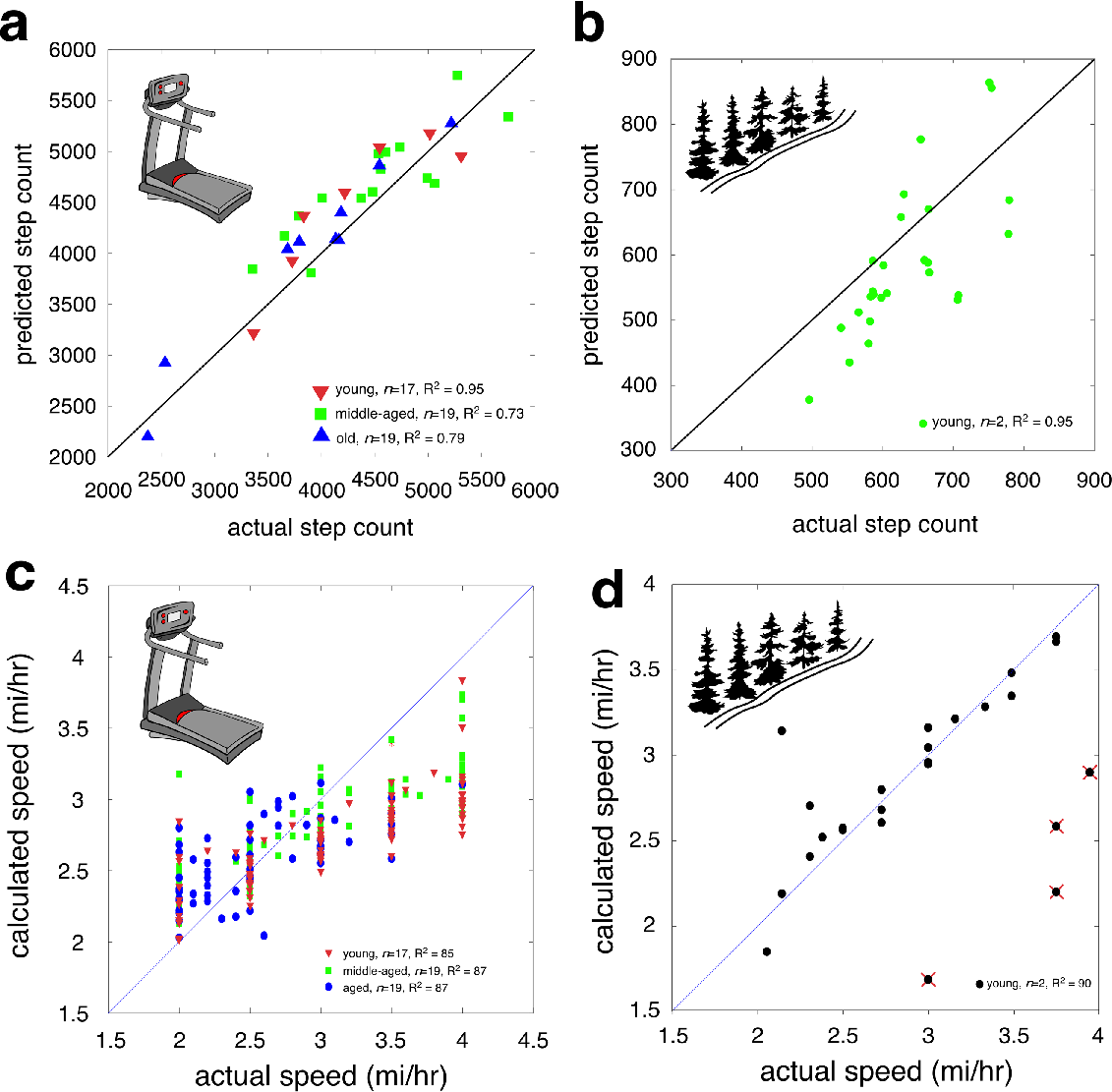
Validation of algorithm to gold standard measures of step count and gait speed under laboratory and naturalistic conditions. **a.** Treadmill locomotion, step count, gold standard video; **b.** Outdoor walking, step count, gold standard manual step count, **c.** Treadmill locomotion, gait speed, gold standard treadmill set speed; **d.** Outdoor walking, gait speed, gold standard distance/time calculation. Four outlier values are depicted; these data points were collected on the same day and time by the same investigator, and likely represent an undetermined error in data collection. They are shown for sake of completeness. Diagonal line depicts performance of perfect algorithm. For treadmill locomotion, three groups assessed including 17 young (blue), 19 middle-aged (green), and 19 old (red) individuals.

We then applied this model to step counts obtained in naturalistic environments, both at the outdoor track and during day-to-day activities (Figure 5b). We observed excellent agreement both for locomotion at the outdoor track (21576 steps predicted, 21867 steps observed, 91.5% average accuracy) and during day-to-day activities (142079 steps predicted, 140461 steps observed, 88.9% average accuracy).

Finally, we applied this model to the data provided by our older, community dwelling adults who kept the cell phone for 24 hours. We note continued excellent agreement of cell-phone derived footsteps compared to manually counted footsteps (77656 steps predicted, 80307 steps observed, 90.15% average accuracy).

*Thresholding algorithm accurately measures gait speed in both laboratory and naturalistic settings.* To test our gait speed predictions, we first examined model performance under the highly controlled situation of treadmill locomotion. As demonstrated in Figure 5c, our model was 86% accurate for 2 mi/hr < gait speed < 4 mi/hr across all age cohorts and gender. For each individual cohort, we noted accuracies of 85% (young), 87% (middle-aged), and 87% (aged).

As shown in this figure, for low gait speeds, our algorithm tends to overestimate gait speeds; conversely, for higher gait speeds, the algorithm tends to underestimate gait speeds. The reason behind this decline of accuracy is suspected to be the change of stride length based on walking frequency [24]. From the figure, it is evident that lower speeds (<2.5 mph) overestimate gait speed due to the higher stride length value; and higher speeds (>3.0 mph) underestimate gait speed due to the lower stride length value. Further analysis suggests that adjusting stride by dominant walking frequency may improve model correlations at both low and high gait speeds.

Finally, we applied this model to data obtained in naturalistic environments at the outdoor track, as provided by our two young subjects (Figure 5d). As shown in the figure, most of the data accurately predicted actual gait speed of two subjects with >90% accuracy.

## Discussion

The above work demonstrates that fast, simple, and well-validated algorithms can accurately and efficiently classify accelerometer data into clinically relevant metrics. These algorithms were validated under a variety of experimental situations, including laboratory environments (that provided extensive control over gait speed and observation conditions), and real-life situations. Our results show that these algorithms report subject activity status with significantly greater precision than that provided by intense subject journaling. Similarly, these algorithms accurately determine subject step count and gait speed for a wide range (both in age and functional status) of clinical subjects. Together, these results suggest that smartphone-derived accelerometer data can provide valuable metrics regarding an individual’s activity status and gait.

There is a rich history regarding algorithmic approaches to analyze activity status and gait speed. Cepstral [25–27], artificial neural network [28–30, among others], hidden Markov model [31–33, among others], support vector machine 34–36, among others] and Bayesian classifier [37–39, among others] approaches have all been validated, with accuracies ranging between 79-97%. There is also a growing literature demonstrating that intrinsic smart phone accelerometers provide valid activity measures in ambulatory human populations [15,40–43, among others]. Here we demonstrate that simple analysis algorithms that do not rely on extensive signal feature extraction or pre-taught learning networks can accurately (86-96%) classify clinically relevant activity and gait metrics from smartphone-derived accelerometer data. Furthermore, we obtained these results with no effort to maintain consistent smartphone orientations. These results add further weight to the concept that smartphones may be repurposed as accurate devices to measure physical activity in a variety of adult, community dwelling populations.

Classification errors are an inevitable outcome of any algorithmic approach, particularly when attempting to separate behaviors whose properties partially overlap. Thus, as we examine activity on a continuum ranging from ‘forgotten phone’ to ‘resting’ to ‘low physical activity’ to ‘high physical activity’, it is unsurprising that we find small errors in classification between two adjacent classes. Our classification errors range from 0.4% (differentiating ‘forgotten phone’ from ‘resting’) to 8% (differentiating ‘resting’ from ‘forgotten phone’ and ‘low physical activity’). As noted above, these accuracies are in the same range as those determined by a variety of algorithmic approaches. One factor potentially responsible for this performance (and consistency to the above-mentioned studies) is that we classify behaviors across narrow time windows, minimizing the effect of activity transitions on overall algorithm performance [44]. Since our classification approach differentiates frequencies within a relatively narrow range (frequencies too low or too high represent unphysiological conditions), our approach is robust to the frequency limitations inherent in windowing, while benefiting from the temporal precision possible by narrow window lengths. These accuracies suggest that subjects forgetting to carry the phone do not degrade our ability to identify resting and low physical activity states. Similarly, our results suggest that our windowed Fourier classification approach can reliably differentiate low from high physical activity states.

Modern pedometers measure step count through a spring-mounted pendulum arm that makes and breaks an electrical circuit (spring-levered), or a beam that deforms a piezoelectric element with each step (piezoelectric). These pedometers are well appreciated to have multiple limitations, including significant step undercounting with slower gait speeds, obese individuals, older adults, or persons with gait impairments [45–48]. Piezoelectric pedometers have higher sensitivity than spring-levered pedometers, and also are not as dependent upon device orientation. However, both pedometers still undercount steps, particularly at lower gait speeds [49]. Our approach of examining acceleration properties across our 8.5 s subwindows yielded an overall accuracy of between 90-92%, including older individuals with known gait problems. Smartphones thus match or best performance of pedometers for measuring step counts. Of note, our algorithms had their least accurate performance determining gait speed. Gait speed is a clinically vital metric [50] with considerable health outcome consequences [51]; yet, most ambulatory studies do not attempt to measure participant gait speed (however, see [52–54]). Inaccuracies associated with our approach may arise from dynamic changes in stride length occurring over different gait speeds.

The ultimate goal of this work is to develop the underlying framework so smartphones contribute accelerometer data as part of a “big data” approach to obtain population-based metrics of human activity. While individual participants may find these metrics personally appealing (by better benchmarking of day to day activities [*e.g.* 55,56]; better health care outcomes [*e.g.* 57,58]; obtaining discounts on health care services, [*e.g.* 59,60]), we envision that the availability of long-term daily measures of physical activity will have significant impact on public healthcare practice and policy [60,61; among others]. Nuanced and rich data assessing physical activity in persons across different demographic, environmental, geographic, and social settings (with privacy concerns addressed, *e.g.* [63]) is absolutely required if policymakers are to have the critical information needed to create or tailor programs to increase population physical activity.

## Acknowledgements

Supported by the Alzheimer’s Association Everyday Technology for Alzheimer’s Care (ETAC) grant 11-206024 (CRH, AKS, SJB), R34MH100460-01 (EHG). The authors would like to thank Samsung USA (R. Cha, C. Sastry) for their technical and material support; P. Moffatt, R.N., Jeanne Costello R.N., and Kelli Kubik R.N. for their help enrolling subjects with functional limitations, and the UNMC EngAge program for their support enrolling functionally intact older adults. AKS thanks Randolph College for material support. Most importantly, we thank our study participants for their generous gifts of time, effort, and interest.

## Abbreviations

MOCA: Montreal Cognitive Assessment

RMS: root mean square

IPA: low physical activity

hPA: high physical activity

FFT: fast Fourier transform

